# Reanalysis of published histological data can help to characterize neuronal death after Spinal Cord Injury

**DOI:** 10.1101/2024.01.22.576608

**Authors:** Pablo Ruiz Amezcua, Nadia Ibáñez Barranco, David Reigada, Irene Novillo Algaba, Altea Soto, M. Asunción Barreda-Manso, Teresa Muñoz- Galdeano, Rodrigo M. Maza, Francisco J. Esteban, Manuel Nieto Díaz

## Abstract

Spinal cord injury (SCI) is a disabling disorder of the spinal cord resulting from trauma or disease. Neuronal death is a central event in the pathophysiology of spinal cord injury. Despite its importance and the large number of research studies carried out, we only have a fragmentary vision of the process focused on the specific targets of each study. It is our opinion that the research community has accumulated enough information which may be reanalyzed with novel tools to get a much more detailed, integrated vision of neuronal death after SCI. This work embeds this vision by creating NeuroCluedo, an open data repository to store and share images as well as the results from their analysis. We have employed this repository to upload the raw and processed images of spinal cord sections from a mouse model of moderate contusive SCI (Reigada et al., 2015) and used this information to: compare manual-, threshold-, and neuronal network-based neuron identifications; and to explore neuronal death at the injury penumbra 21 days after injury and the neuroprotective effects of the anti-apoptotic drug ucf-101. The results from these analyses i) indicate that the three identification methods yield coherent estimates of the total number of neurons per section; ii) identified the neural network as the optimal method, even in spinal sections with major artifacts and marked autofluorescence associated with spinal damage; iii) characterize neuronal distribution among Rexed laminae in the mice T11; iv) reveal that neuronal death distributes through all the gray matter in the penumbrae sections closer to the injury epicenter but concentrate in the intermediate region in sections located farther away; and that v) antiapoptotic effects of UCF-101 are highest in the intermediate region of the gray substance of the caudal segments closest to the injury epicenter. All methods and results, including raw and processed images, software, macros, and scripts, together with all data matrixes and results have been deposited and documented in the Open Science Framework (OSF) repository Neurocluedo (https://osf.io/n32z9/).

## INTRODUCTION

According to the WHO (Bickenbach *et al*., 2013), the term Spinal Cord Injury (SCI) refers to the damage to the spinal cord resulting from either external physical impact (traumatic SCI) or from disease and degeneration (non-traumatic SCI). Although there is no reliable estimate of global prevalence, up to 7 million SCI patients in the world (Singh *et al.,* 2014) live with permanent disabilities of enormous personal, medical, social, and economic costs ($1.2 to 5 million/patient in the USA according to NSCISC, 2021). SCI causes extensive alterations in the Central Nervous System (CNS) leading to impaired motor, sensory, and autonomic functions. Research efforts devoted during the past 40 years have yielded multiple therapies, several of which have been tested in clinical trials (Dietz *et al.,* 2022). However, to date, only early surgery interventions, intensive care management, and rehabilitation remain the cornerstones for SCI treatment (Ahuja *et al*., 2017; Donovan & Kirshblum, 2018).

Trauma to the spinal cord triggers a complex pathophysiology that extends the initial damage to distant regions of the central nervous system during the following weeks and months (Ahuja *et al*. 2017). Cellular death, particularly which of neurons, is a central event of SCI pathophysiology and a major determinant of the resulting functional deficits. In traumatic SCI, neuronal death is biphasic. Initially necrotic, driven by the physical breakage of axons and somas, the subsequent alteration of the spinal cord environment and pathophysiological events trigger secondary waves of neuronal death that extend throughout the spinal cord and the brain (Hassannejad *et al*., 2018). Contrary to the necrotic death that characterizes the primary damage, secondary damage is characterized by programmed processes (Kuzhandaivel *et al*., 2011; Zhang *et al*., 2012; Liu *et al*., 2015; Wu & Lipinski, 2019) which may be regulated with appropriate neuroprotective treatments (Zhang *et al*., 2012).

Histological analyses back in the 1990s and the early 2000s provided spatial and temporal characterizations of neuronal death in animal models of spinal cord injury, describing the onset and extension of necrotic and apoptotic processes (see, for example, Beattie *et al.,* 2002). Since then, the increase in knowledge, technologies, and techniques has allowed identifying novel forms of PCD acting on the injured spinal cord as well as precisely characterizing the effects of neuroprotective treatments (for review, see Shi *et al.,* 2021 and references therein). The recent development of single-cell/single-nucleus transcriptomics has opened new opportunities to analyze in detail the different processes affecting neurons with single-cell resolution (Sathyamurthy *et al.,* 2018; Matson *et al*., 2022). Integration of all this information will provide a detailed and precise view of the temporal and spatial course of neuronal death after SCI, contributing to determining its causes, mechanism, and involved processes, as well as to better understand the effects of neuroprotective treatments. However, this integration is lacking and, to our knowledge, no comprehensive characterizations of neuronal death have been published since the studies from 20-30 years ago. Here, we hypothesize that the histological information gathered through the years can be reanalyzed by employing novel techniques to fill this gap. In this respect, this work tries to contribute by:

1. creating an open data repository to store and share images as well as the results from their analysis;
2. depositing and documenting 168 images of full transversal sections of mice spinal cords from the study by Reigada *et al*. (2015) on the neuroprotective effects of Ucf-101 in the damaged spinal cord, and using the stored information to:
3. compare the precision and reproducibility of manual-, threshold- and neuronal network-based neuron identification methods; and
4. explore how neuronal death distributes within the spinal cord sections in Reigada’s mouse model of SCI and which neurons become protected by the treatment with the anti-apoptotic drug ucf-101.

## MATERIAL AND METHODS

### 1. Data

#### 1.1. Open Access Repository

We have employed Open Science Framework (https://osf.io; Foster & Deardorff, 2017) to store and share all information from this study.

#### 1.2. Images and associated data

The dataset comprises 168 confocal images from full cross-sections of female C57BL/6J mice spinal cords up to 1.5 mm rostral and caudal to the injury epicenter. Images correspond to sections of spinal cords from 2 sham (laminectomy only) and 11 injured (50 Kdynes moderate contusion at spinal segment T11 using an Infinity Horizon Impactor) individuals (5 treated with ucf-101 and 6 with vehicle). These images were originally employed to analyze the neuroprotective effects of the drug ucf-101 (Reigada *et al*., 2015). Injured and sham animals were euthanized 21 days after surgery and, after processing, their spinal cords were cut into 20µm thick transversal sections. For neuronal quantification, sections were stained with an antibody against the neuronal marker NeuN (Millipore cat#MAB377, RRID: AB_11210778) and the nuclei marker DAPI (4′,6-diamidino-2-phenylindole, Sigma Aldrich) and imaged with a Leica TCS SP5 fast-scan confocal microscope equipped with a PL APO 20x/0.70 CS 20X objective.

Additional information can be accessed in the original publication (Reigada *et al*., 2015) and at NeuroCLUEDO (https://osf.io/qe9ys/). The repository includes confocal images in .lif format plus metadata for each individual. Each .lif file contains all imaged sections from an individual. Metadata comprises MIASCI data as well as information on each animal, section, and image contained in the .lif files.

For all analyses in this study, we have transformed the confocal images from each individual stored in each .lif file into independent .tif files. The transformation was carried out using FIJI (ImageJ vs 1.53c; Schneider *et al.,* 2012) and involved maximum intensity projection of the focal planes of the confocal image and its conversion into RGB format.

### 2. Experimental Design

Three identification methods were compared, one based on manual identifications, another based on semi-automatic thresholding, and the last based on a deep learning neuronal network. The experimental design is summarized in Figure 1. 20 images from those stored in the NeuroCLUEDO repository were employed to compare neuron identification methods. Both the total number of identified neurons and the position of each identified neuron were employed in these comparisons. Manual quantifications were employed to obtain a consensus estimate of the number of spinal neurons in each section of the comparison set.

**Figure 1.**
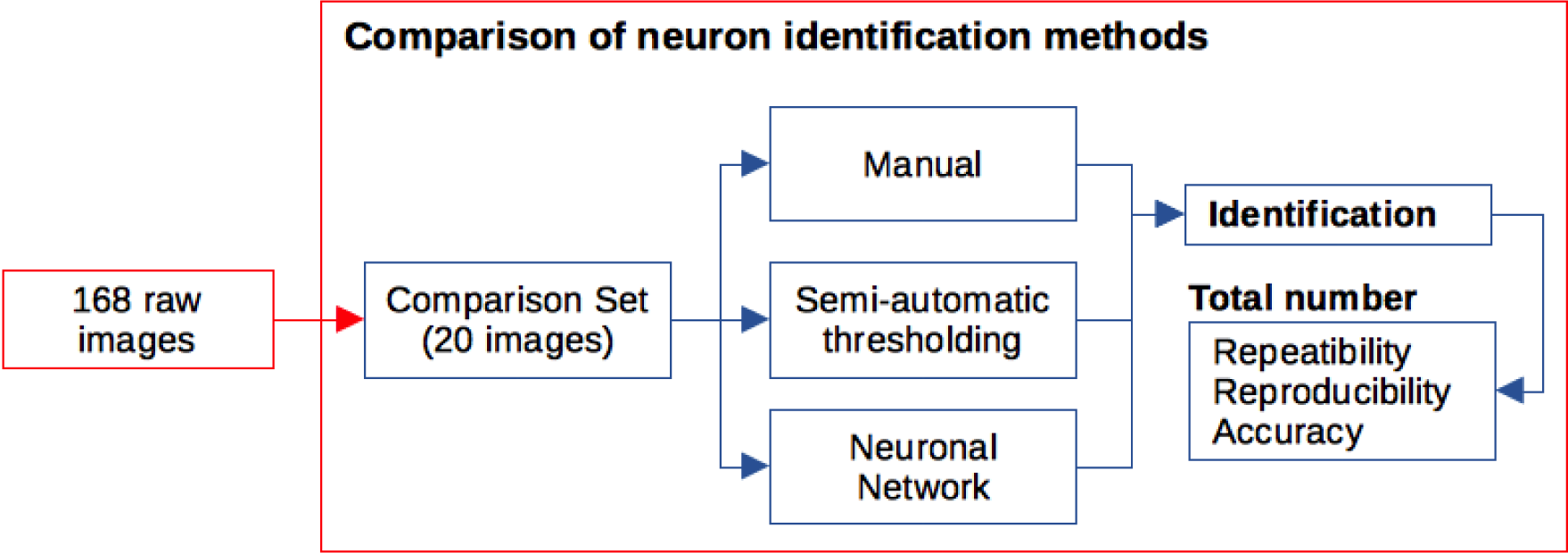
Summary of the study design. 168 images of full spinal cord sections from 11 sham, injured, and treated mice were deposited at OSF. 20 of these images were selected as the comparison set. Data on the identified neurons and their total number per section were employed to compare manual, threshold-based, and neuronal network-based methods of neuronal identification.

### 3. Neuron identification methods

The comparison set of images was analyzed using three different methods: (i) a manual analysis carried out by six observers using the Cell Counter plugin from Fiji; (ii) a threshold-based identification with a custom Fiji macro; and (iii) a neural network-based identification employing the deep-learning-based image analysis approach TruAI integrated CellSens Dimensions software (Olympus). For details of the three methods, see supplementary materials.

### 4. Comparative analysis of neural identifications

The neurons identified by each method were compared by overlapping their positions or segmentations obtained with each method for each image, and repeatability, reproducibility and accuracy of neuronal quantifications were estimated. For details, see Supplementary materials.

### 5. Reanalysis of Ucf-101 effects on neuronal survival

42 sections were included to reanalyze the effect of Ucf-101 on neuronal survival. Neurons were identified using the neural network-based identification above indicated. With the aim to obtain an estimation of the neurons present in each lamina and nucleus comprised in the Mouse Spinal Cord section of the Allen Brain Atlas, images from each section under analysis were registered to a reference map of the Rexed laminae in thoracic segment 11 obtained from https://mousespinal.brain-map.org/imageseries/showref.html (last date accessed august 2023). For details, see supplementary materials.

## RESULTS

### A. REPOSITORY

Following the first objective of this study, we have created an open-access project within the OSF repository where all images, data, and results (see table below) from this article are freely available under the Creative Commons (CCO 1.0 Universal) license. The project, named NeuroCLUEDO|Spinal Cord Injury (https://osf.io/n32z9/), is aimed to determine which, where, and how neurons die after SCI. It is structured in 3 major components:

- a documented **image repository** of spinal cord sections stained with neuronal markers.
- **test benches** for comparing methods for image analysis in the naïve and injured spinal cords.
- **results** from the analysis of the images in NeuroCLUEDO repository.

All raw and processed images together with any additional information described here and the results from the comparison of neuronal detection methods and the reanalysis of the effects of ucf-101 on neuronal survival after SCI have been distributed among the three components of NeuroCLUEDO as detailed in Table 1.

**Table 1.**
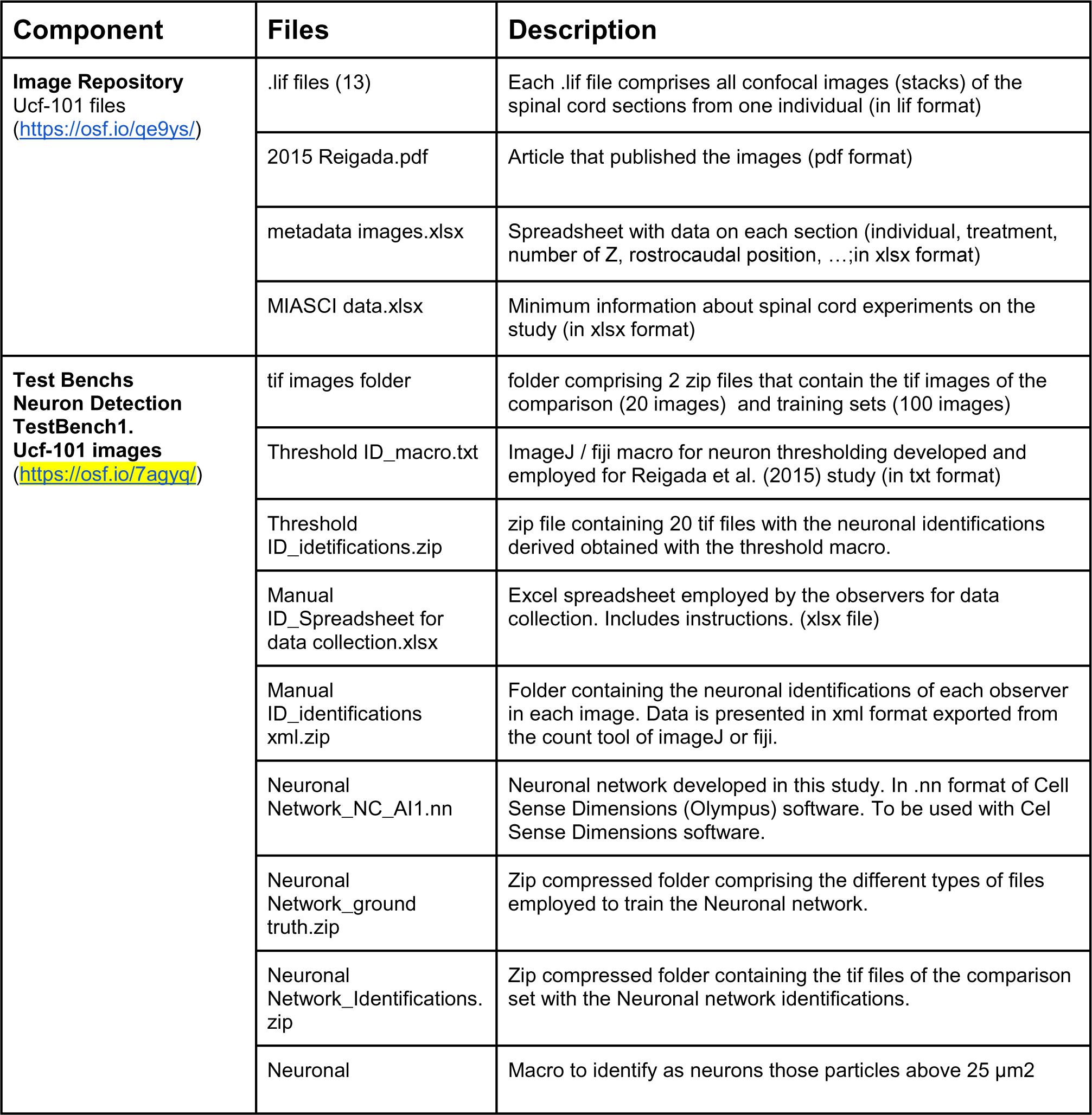
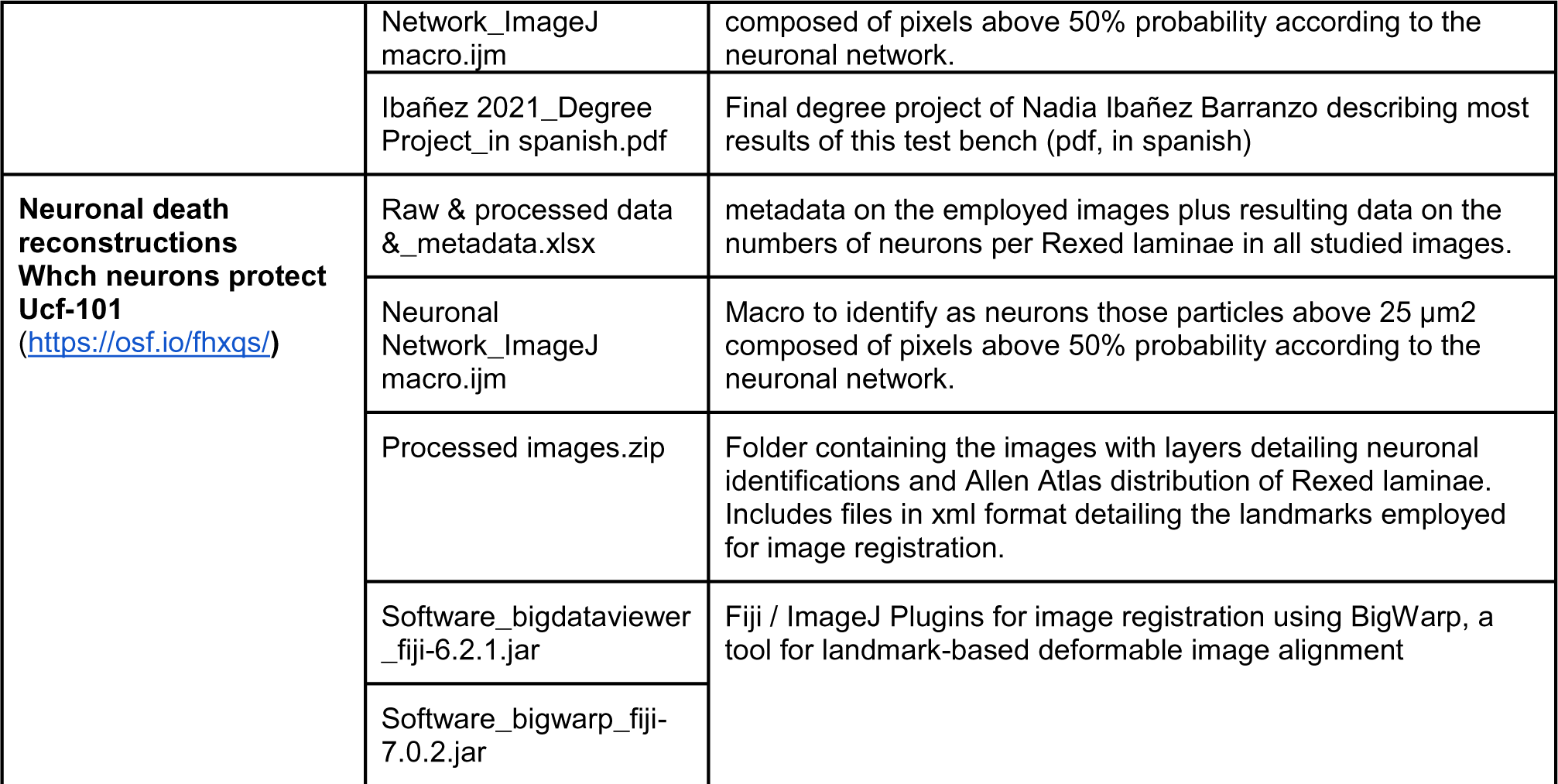
Contents related to this study in OSF repository (. https://osf.io/7agyq/). The table lists and describes all files related to this study present in each of the three components of NeuroCLUEDO. Additional contents related to other analyses can be present in each component within specific subcomponents. Each component and subcomponent has its own wiki page with additional details and information.

### B. COMPARISON OF METHODS FOR IDENTIFYING NEURONS

#### B.1. Neuronal identifications

In this study we have compared the neurons identified through manual, threshold-based, and neuronal network-based methods in 20 spinal cord sections originally studied in Reigada *et al*. (2015). All manual identifications, together with threshold- and neuronal network-based identifications, can be accessed at NeuroCluedo repository (https://osf.io/7agyq/).

As shown in Figure 2, each method yields different outputs. Manual identifications result in the X and Y coordinates of each neuron, thresholding results in a binary segmentation of the image, whereas Neuronal Network assigns a probability of being part of a neuronal nucleus to each pixel in the image. Despite these differences, the three methods yield broadly comparable identifications, all within the gray matter –an established condition in manual and threshold-based identifications but not in Neuronal network-based ones–. In addition, the Neuronal network was able to exclude most artifacts as did manual identifications whereas, on the contrary, the threshold method included multiple artifacts.

**Figure 2:**
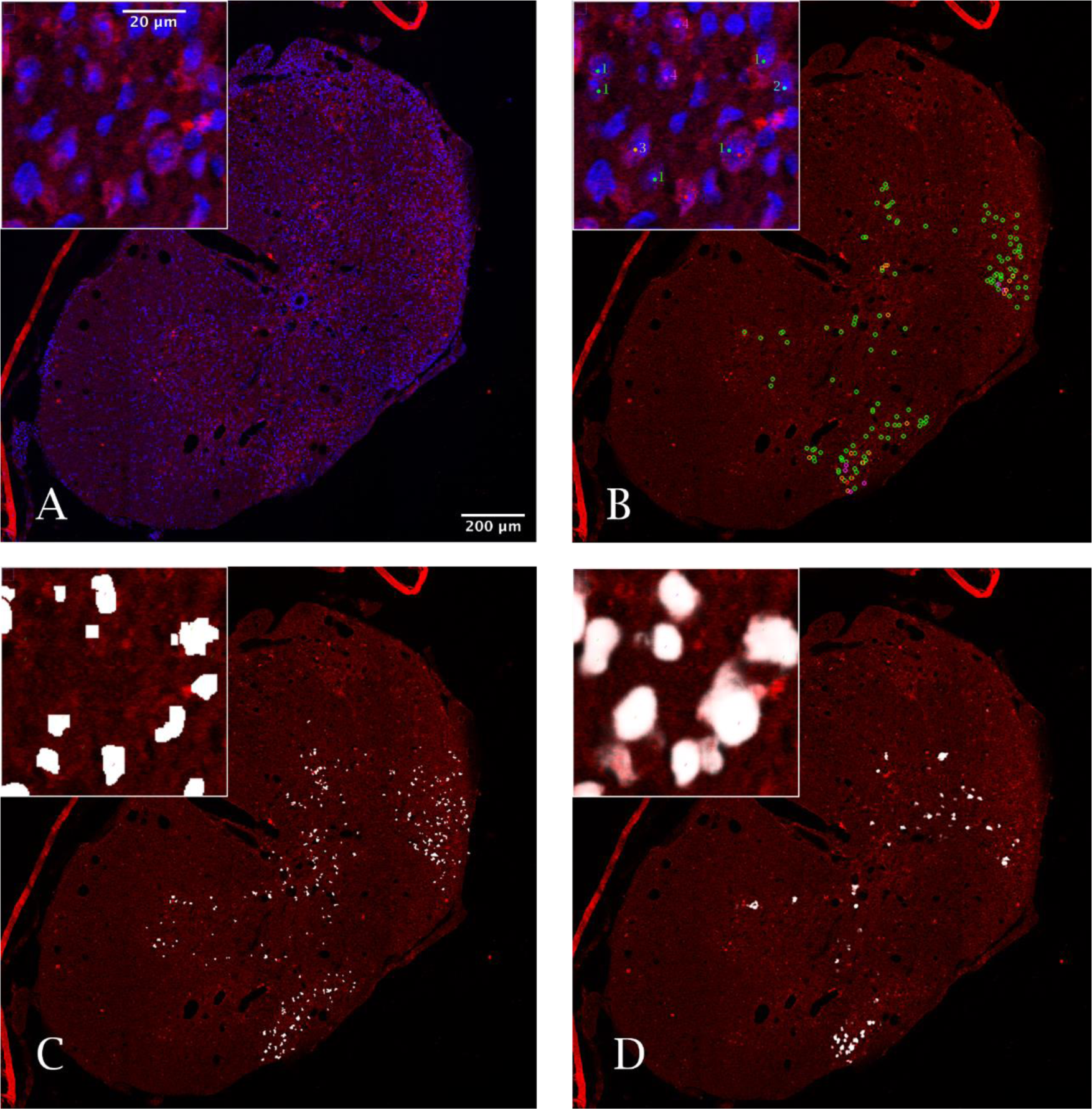
Neuron identifications in the spinal cord. The figure shows the identifications obtained with the three methods in the full section and a detail (insert) of image 188. A. Image under analysis. B. Manual identifications. Numbers represent the number of times each neuron was identified after 5 analyses. C. Segmentations obtained with the threshold-based method. D. Probabilities of each pixel to be part of a neuron (in a 0 to 255 intensity scale) according to the neuronal network.

To compare the results and assess the advantages and disadvantages of each procedure, we first overlapped the neuronal identifications of each image obtained with each method and observer to explore the coherence in the identifications of each neuron. In the case of the manual method, 3 or 5 analyses (from 2 or 3 observers, respectively) *per* image were overlapped and each neuron was characterized by the number of analyses in which it was identified. Most neurons were identified in a single analysis and can therefore be considered highly doubtful (see Table S1 at supplementary materials). These doubtful neurons are particularly abundant in sections of untreated injured cords. Neurons identified by all analysts, which can be considered undoubtfully as neurons, are also abundant, particularly in sections from control individuals. Neurons identified in several, but not all, analyses represent a minority in most sections (Table S3).

To compare with the identifications of the threshold-based method and the neural network, their identifications were overlapped on the manual identifications in five images (Figure 3 and Table S4). Both systems identified most neurons identified in 3, 4, or 5 out of the 5 manual analyses. Conversely, those neurons with less agreement among manual identifications (1 or 2 out of 5) showed a markedly lesser agreement with the macro and IA identifications suggesting that assignment to neurons is at least doubtful. These results suggest that both the macro and the neural network identify as neurons the cells with the highest probability of belonging to this cell type and only show greater discrepancies with the human analysts in the doubtful cases. On the other hand, both the macro and the neural network identified neurons unidentified by the analysts (value 0 in Figure 3 and Table S4). The number of these neurons absent in the manual identifications is higher in the macro results (up to 87% additional cells) than in the case of the neural network (19%). In most cases, these cells do not correspond to neurons or their assignment is very doubtful. Therefore, it can be deduced that the neural network is capable of quite effectively discriminating cells with high labeling due to autofluorescence or artifacts that do not correspond to neurons, while the macro network has difficulties in this aspect.

**Figure 3.**
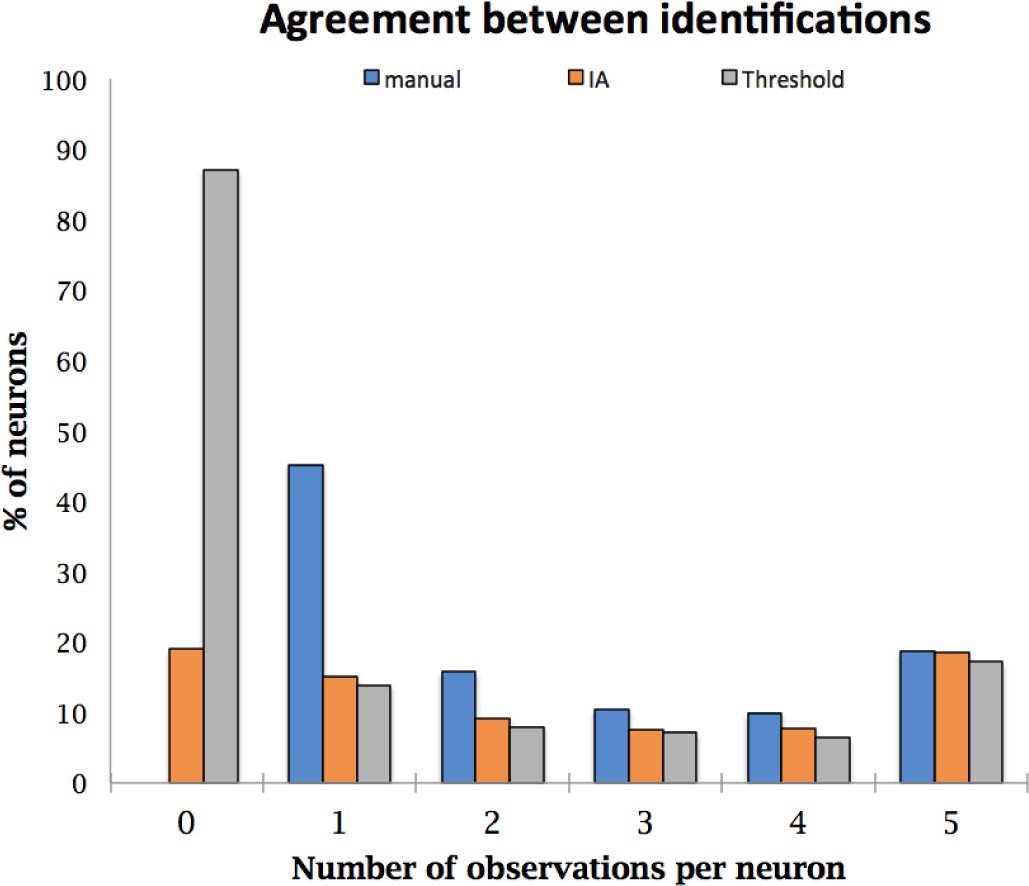
Agreement of manual, threshold and IA based identifications. The graphic summarizes the data shown in Table 4. Data is relativized to the total number of neurons identified manually.

#### B.2. Number of neurons

The number of neurons in a region or section is one of the parameters that can be derived from the identifications that is most used in scientific studies of SCI and other pathologies of the nervous system. The neuron counts derived from each observer (along with the time spent) are summarized in Table S5.

##### B.2.1. Repeatability

The repeatability (Figure 4) that an individual presents with respect to himself was studied by comparing the two values that a subject has recorded from the same spinal section. To do this, the difference of both values was represented against their average. These studies, like the subsequent reproducibility studies, have been carried out only on manual quantifications because, as mentioned in the materials and methods section, both the threshold method and the neural network are 100% repeatable and reproducible (although there are some manual steps of the threshold method that could incorporate variability that has not been considered in this work). The Bland and Altman graphs (Figure 4) show important changes in the number of neurons estimated in the repeated measurements throughout the entire range of values measured (from null values to values of 600 neurons). Although most of the differences between measurements repeated by the same observer are concentrated around ± 50 (95% confidence intervals +111.71; -137 .51).

**Figure 4.**
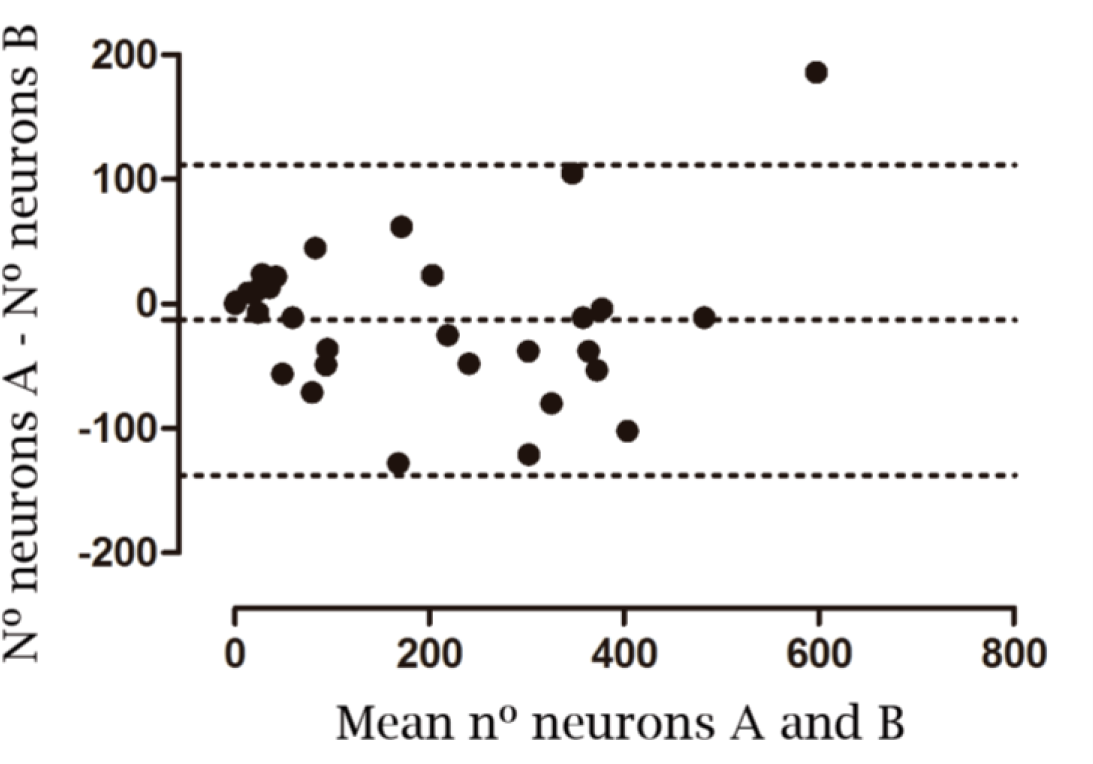
Repeatability of manual identifications. Bland-Altman graphic representing repeatability of replicate identification by observers. The graph compares the differences between repeated counts of total number of neurons from the same spinal cord section versus their mean. The horizontal lines refer to the mean values (n=-12.9) ± the 95% confidence intervals.

Various factors can influence repeatability. In this work, three of them have been considered: number of neurons, experience of the volunteers and state of the spinal section. To assess its influence, linear models were adjusted relating the difference between the repeated counts (inverse of repeatability) in a section against the average number of neurons counted, the experience of the analyst (differentiating between beginners, advanced and experts) and the condition of the section (differentiating sections from uninjured spines, sections near the injury epicenter, and sections distant from the epicenter). The comparison of the fitting of the resulting models (see supplementary material online https://osf.io/7agyq/l) using analysis of variance (ANOVA) showed that the best fit (p=0.039) is achieved by the model that relates the range to the mean number of neurons and the experience of the observer (Adjusted R^2^=0.500, p<0.001).

##### B.2.2. Reproducibility

Reproducibility between observers was studied by comparing the mean values of the different sections among analysts. As can be seen in the Bland-Altman graph (Figure 5), reproducibility is similar to repeatability, only with a slight increase in the 95% confidence intervals (+151.36;-159.24).

**Figure 5.**
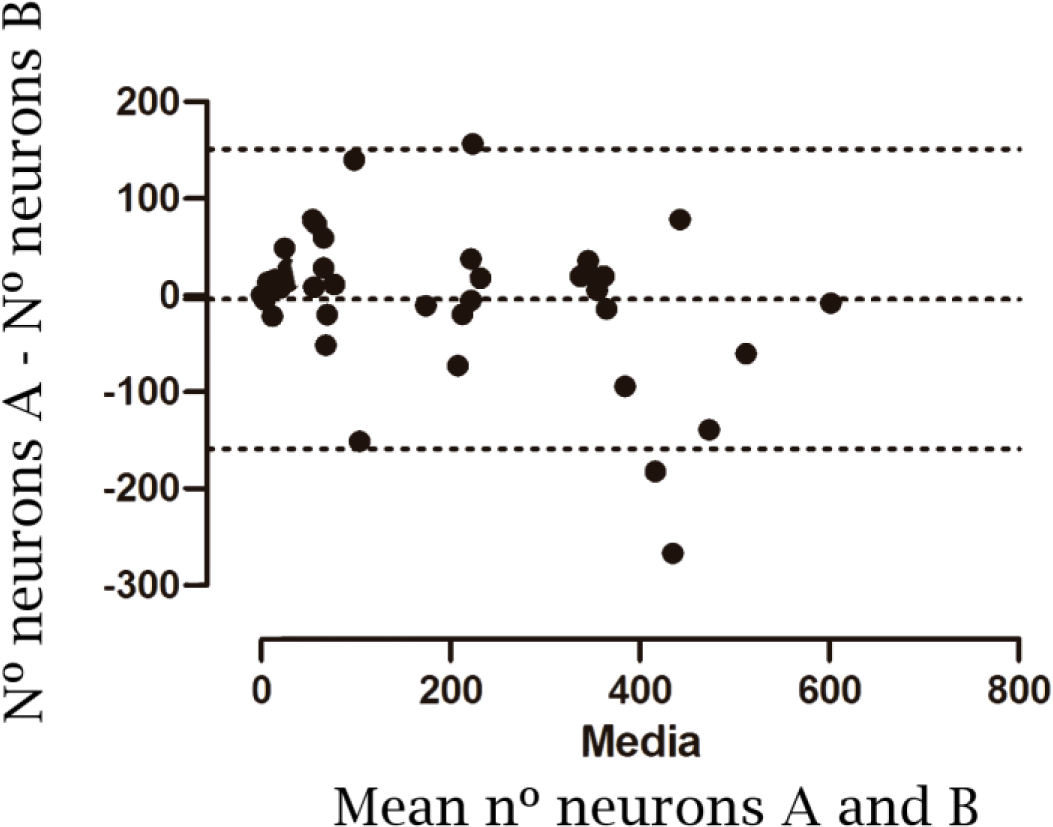
Reproducibility of manual identifications. Graphic representation of reproducibility according to the Bland-Altman graph. The horizontal lines refer to the mean values (n= -3.93) ± the 95% confidence intervals.

As in the repeatability tests, the effect of the number of neurons, analyst experience, and the type of section (non-injured, near and distant from injury epicenter) on reproducibility was assessed by again comparing linear models using ANOVA. In this case, none of the factors has a significant effect. Despite the fact that the model that only considers the number of neurons has a very limited and insignificant fit (adjusted R² = 0.113, p = 0.0193) it is the best model of the three (see supplementary material online https://osf.io/7agyq/l).

##### B.2.3. Accuracy

Accuracy of the manual, threshold-based and Artificial Intelligence-based neuronal counts was estimated by comparing the obtained values against the Reference Number of Neurons (RNN) derived from the weighted consensus of the manual estimations for each image (Figure 6 and Table S6).

**Figure 6.**
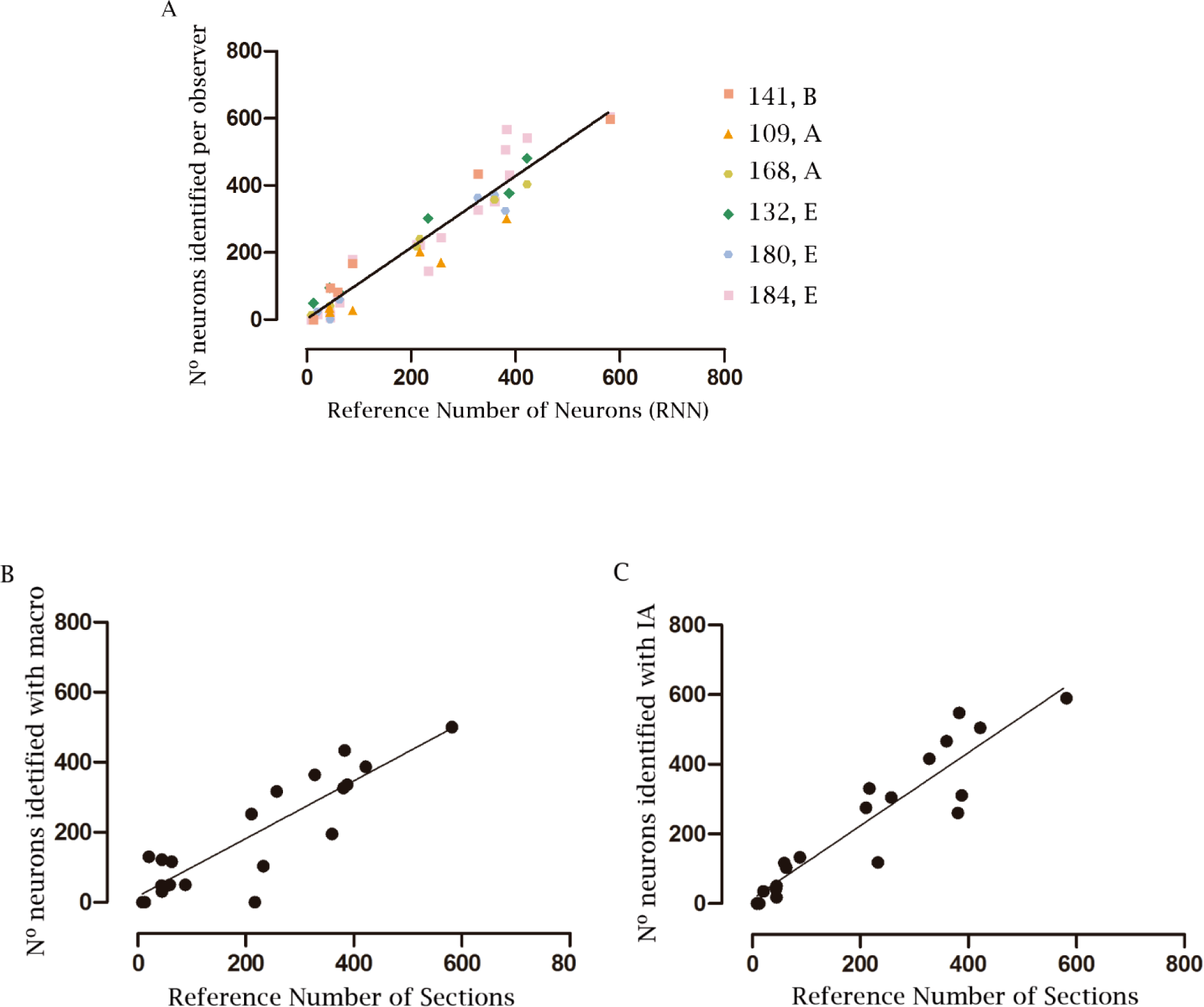
Accuracy of neuronal counts from manual, threshold-based and neuronal network-based identifications. Comparison of the RNN with the estimated values in each manual observation (A), threshold-based identifications (B), and Neuronal Network-based identifications (C). In A, the symbols and colors in A indicate each of the observers, whereas B, A, and E code for beginner, advanced, and expert, respectively.

As shown in figure 6, manual counts show a high correlation (Pearson’s correlation coefficient = 0.9237, p<0.001). Disparity with respect to the RNN concentrates on the identifications by volunteers 141 (beginner) and 184 (beginner aided by an expert). The threshold- and neuronal network-based methods also result in values highly correlated with the RNN. However, the threshold-based method presents a lower correlation (correlation coefficient = 0.7779, p<0.001) compared to the neuronal network identifications (correlation coefficient = 0.8636, p<0.001). Despite this difference, the correlation between the number of neurons estimated by the threshold method and the RNN is highly significant (see Figure 7), so it can be considered that the values obtained in the article by Reigada *et al*. (2015) using this threshold method are an acceptable estimate.

**Figure 7.**
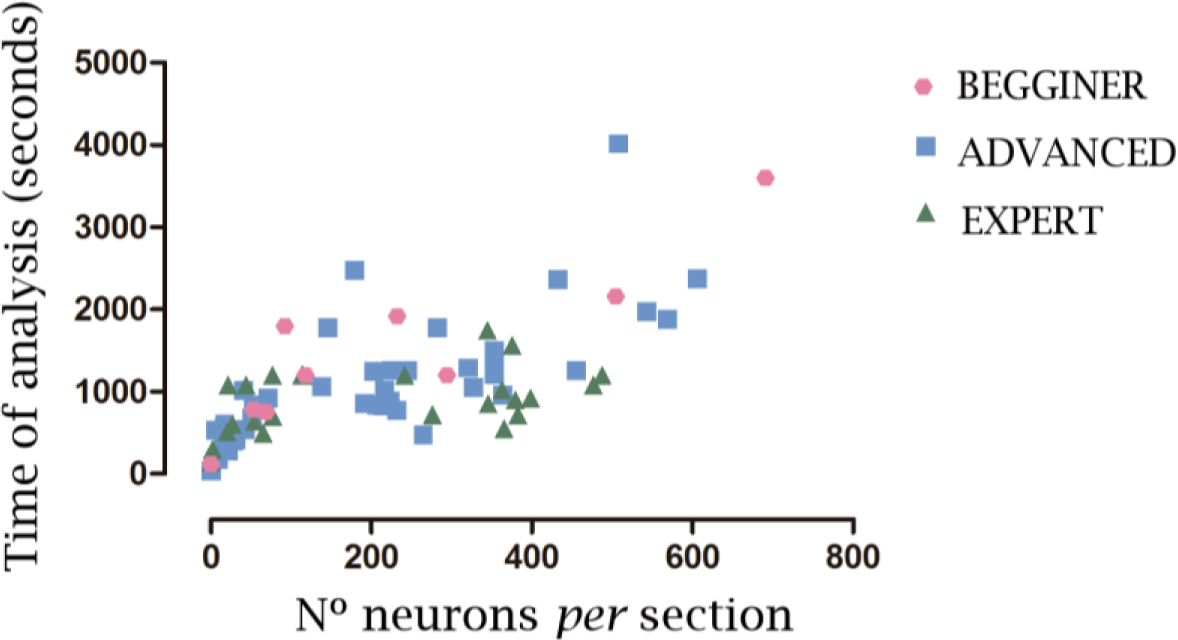
Time of analysis. Relationship between the times to analyze an image versus the number of neurons identified in the image and the observer’s experience. The symbols and colors on the graph correspond to the experience of the observers.

#### B.3. Effort: processing time and experience

The time of analysis was incorporated in this study as a measure of the effort made to identify the neurons in an image. For manual identifications, the average time required to analyze an image has been 17 minutes and 40 seconds with a standard deviation of 12 minutes 03 seconds and a range from 30 seconds to 67 minutes 0 seconds (Table S5). As for reproducibility and repeatability, 3 variables can explain the differences in the time spent: the number of neurons present in the image, their condition, and the experience of the analyst. To assess the effect of these possible factors, linear models were used, incorporating the variables step by step (see supplementary material online https://osf.io/snjwk/). According to ANOVA, the best fit is achieved by a model comprising the three predictive variables although improvement is barely significant (p=0.026) compared to a model only considering the number of neurons and the observer experience (p<0.001). Moreover, this second model is able to explain a slightly more variability (adjusted R^2^=0.585) than the model involving the 3 variables (adjusted R^2^=0.584). Therefore, the time necessary to manually analyze an image increases significantly with the number of neurons in the image, with an estimated 3.26 seconds per neuron, except in the identifications of expert observers whose time per neuron decreases significantly (p=0.002) to less than 1 second per neuron.

On the other hand, the time spent in the threshold-based identifications was estimated at 3-4 minutes per image while, using the neural network developed in this work, the identifications of the 20 images required 3 minutes in total. In both cases, the time does not vary depending on the experience of the observer or the characteristics of the section, but it can be influenced by the processing speed of the computer equipment. In both cases, the time to develop the neuronal network and the macro for threshold-based identifications should be added to the processing time. In the case of the neural network, it was necessary to invest 15 hours of work for training and preparation, whereas the design and preparation of the threshold-based macro required more than 30 hours of work.

### C. EFFECTS OF INJURY AND UCF-101 TREATMENT

#### C.1. General description of the obtained data

In this second part of the study, we compared the effects of SCI with or without treatment with ucf-101 on neuronal death and its spatial distribution. We focussed our analysis on the sections 0.6 to 1.2 mm caudal to the injury epicenter, where Reigada *et al*. (2015) observed a significant effect of ucf-101 on neuronal survival. The images from 43 sections (see Metadata images.xlsx file in https://osf.io/qe9ys/ for details) were processed to identify and quantify the number of neurons in each Rexed laminae. Alterations of the sections, staining failures, and other problems precluded the analysis in 4 of these sections, therefore, 39 sections were included in these analyses (see Table S7). The final data matrix describing the number of neurons in each Rexed laminae *per* section is available at OSF (https://osf.io/brq5k).

A median of 283 neurons were recorded in 4 spinal cord sections from undamaged animals. As shown in Figure 8 and Table S8, image registry allowed determining the distribution of neurons among Rexed laminae showing the highest numbers of neurons in dorsal laminae, particularly in L3 and L4, as well as in the lower part of lamina 7. On the contrary, minimal counts are detected in laminae 9 as well as in the small nuclei (ICI, IML, or D). To evaluate the congruence of image registration, we compared the number of neurons in each laminae obtained here with the numbers derived from the registered image of the T11 segment of P56 days old mouse from the reference atlas for the mouse spinal cord available at the Allen Brain Atlas (https://mousespinal.brain-map.org/imageseries/showref.html). As shown in Table 8, the percentages of neurons per lamina obtained in our analyses show good agreement with those derived from the Allen atlas, with only the values from L1, L5L, and IMM shown discrepancies above a 2 fold. Conversely, the absolute number of neurons *per* lamina is highly divergent as could be expected considering the differences in tissue processing, staining, and imaging.

**Figure 8:**
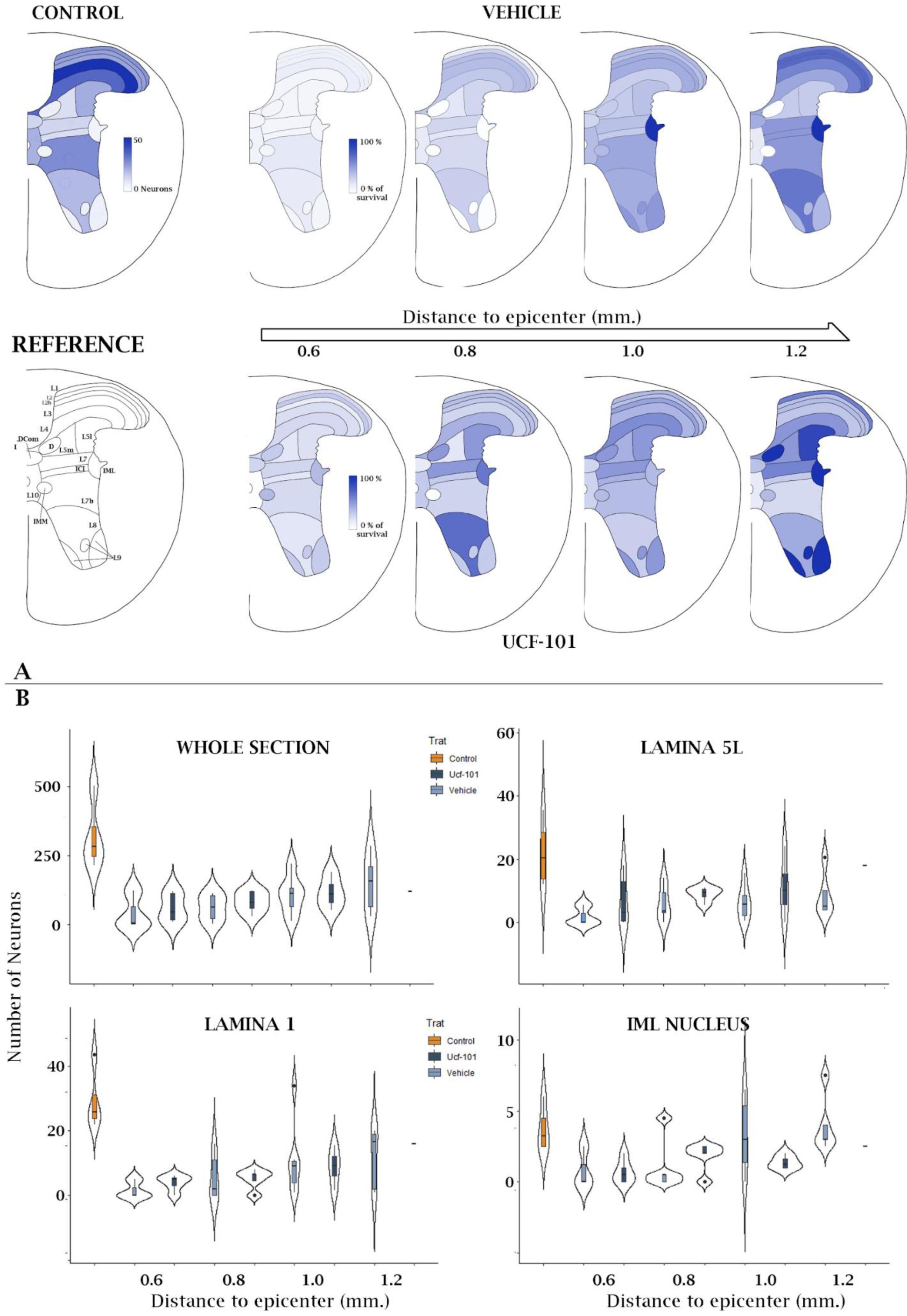
Changes in the number of neurons after SCI and ucf-101 treatment. **A**. The spinal cord drawings illustrate the number of neurons in each Rexed lamina of the T11 segment of the naïve spinal cord (Control) and the percentage of surviving neurons (relative to naïve spinal cord) in each lamina at 0.6, 0.8, 1.0, and 1. 2 millimeters caudal to the lesion epicenter after SCI plus vehicle (above) and ucf-101 treatment (below). Also illustrated is the identification of each Rexed lamina according to Allen’s mouse spinal cord atlas (https://mousespinal.brain-map.org/imageseries/showref.html). Data correspond to the median values of the different individuals and sections analyses available in table 9. The spine image along with the identification of Rexed slices was modified from the reference atlas of the Allen mouse spinal cord atlas. **B.** Violin plots depicting the number of neurons in the whole section and in selected laminae of un-damaged and injured spinal cords. Violin plots also represent the median (red circle) and the data points (black circles). L5L and L1 laminae are included as representative of laminae with high and low effects of ucf-101 on neuronal survival, respectively. IML nucleus is also included to illustrate the anomalous values observed in some laminae and nucleus due to limitations in their identification during image registration. Violin plots for all other laminae are available as supplementary material.

#### C.2. Neuronal loss after SCI

To characterize the neuronal losses 21 days after injury in the present mouse model, we compared the neuronal counts in sections from un-injured individuals with the counts of sections 0.6 to 1.2 mm caudal to the injury epicenter from 6 individuals that underwent a moderate contusion (50 Kdynes) at spinal segment T11. Compared to the median of 283 neurons recorded in the sections of the un-injured individuals, the sections from injured individuals show a reduction in the number of neurons in all laminae at the 4 distances under study. Neuronal loss is massive in the sections 0.6 mm away from the epicenter (median total number of neurons=6.5, ∼2.3% survival) and gradually recovers with a pace of nearly 20% neurons every 200 microns to reach a 56% survival rate at 1.2 mm (see Table S9). Gradual increase of surviving neurons is observed in all laminae with some degree of heterogeneity. The most dorsal laminae (L1, L2, and L2b) show high neuronal survival, particularly at 1.2 mm away from the injury (reaching 70% of control values). High neuronal survival is also observed in L7, L9, and the intermediolateral nucleus (IML), the last two with high survival values already at 1 mm away from the epicenter although the lower number of neurons recorded in these regions may bias the results. Conversely, survival is particularly poor in laminae L5L, L5M, L8, and L10, as well as in nuclei D, ICI, and LDCom, all in the intermediate gray substance. Neuronal survival at 1.2 mm away from the epicenter is below the 40% in all these regions and, in some cases (D nucleus), even below the 20% of control numbers. Kruskal Wallis analyses (shown in Table S9) confirm the highly significant losses of neurons in all laminae from sections sampled 0.6 and 0.8 mm caudal to injury, whereas at 1 and particularly 1.2, losses are marginal or non-significant. Despite general trends, variability among individuals within the same condition is high (see violin plots in Figure 8), which may preclude statistical significance.

#### C.3. Neurons protected by Ucf-101

Analyses from Reigada *et al* (2015) revealed that treatment with ucf-101 results in a localized neuroprotection, centered in the penumbrae segments 0.6 to 1.2 mm caudal to the injury epicenter. Focusing on these sections, our results confirm ucf-101 neuroprotection in the whole sections in this region; however, contrary to Reigada *et al*. (2015), protection appears restricted to the sections closer the injury epicenter (0.6 and 0.8 mm caudal to the injury epicenter see Table 9) while the numbers of surviving neurons become even at 1 mm caudal to the epicenter. Data from 1.2 mm is too sparse in the ucf-101 group to draw any significant observations. A deeper exploration of the distribution of ucf-101 neuroprotection among the neurons in the different Rexed laminae and spinal cord nucleus (Table 9 and Figure 9) reveals that ucf-101 neuroprotection is heterogeneous, and appears to concentrate in specific laminae and nucleus of the intermediate region (laminae 5L, 7, 10, D, IML, LDCom), interestingly leaving out lamina 5M. These Ucf-101 protective effects appear to be consistent across the examined distances to the injury epicenter whereas other effects such as the protection in laminae 1, 2, 8, and 9 are only observed at specific distances (Figure 9).

## DISCUSSION

Neuronal death following SCI is a complex and multifaceted process. Research suggests that different types of spinal neurons may indeed exhibit varying susceptibilities to secondary degeneration. While there is evidence to support these differences (Hassannejad *et al*., 2018), it is important to note that the exact mechanisms and extent of susceptibility can be influenced by various factors, including the location and severity of the injury, as well as the specific animal model used for research.

Recent technological breakthroughs –including image analysis and -omics technologies with single-cell resolution, tools for processing big data, and availability of repositories capable of storing and sharing gigabytes of information– promise to revolutionize Biomedicine (Luo *et al*., 2016). The present study stems from this background to develop a framework to integrate histological and -omics information and tools for the analysis of neuronal death after SCI. Here we focus on histological images of SCI already available in the laboratories and which may be a key resource for the analysis of SCI. To this aim, we have created an open-access repository that stores 168 images of transversal sections of mice spinal cords from the study by Reigada et al. (2015) on the neuroprotective effects of ucf-101 in the damaged spinal cord. The images have been employed to demonstrate the outperformance of neuronal networks for neuron identification in histological preparations of damaged spinal cord; and to explore how neuronal death is spatially distributed in a mouse model of contusive SCI and which neurons become protected after treatment with ucf-101 anti-apoptotic drug.

### Repository on spinal cord injury

The primary contribution of this study is the establishment of an open-access repository, known as NeuroCluedo | Spinal Cord Injury, hosted on the Open Science Framework (OSF). This repository contains a comprehensive collection of histological images obtained from animal models of SCI, documented in accordance with the MIASCI standard (Lemmon *et al*., 2014). To the best of our knowledge, NeuroCluedo represents the first open-access repository of histological images from animal models of SCI. It encompasses full transverse sections from 13 individuals with and without SCI, and we are committed to continuously expanding this repository with additional individuals.

Moreover, NeuroCluedo serves as a comprehensive archival resource, housing not only the raw images but also all processed images, data, and results derived from image analyses. Additionally, it provides access to the protocols, scripts, and tools employed, all of which are readily available for exploration, utilization, or comparison by the research community. While the current contents of the NeuroCluedo repository primarily reflect the outcomes of this study, our intention is to broaden its scope by incorporating images from other investigations and facilitating comparisons of various methodologies. This expansion encompasses aspects ranging from staining techniques to image acquisition and registration processes, and aims to contribute to our understanding of neuronal death following SCI, the effects of treatments, and the vulnerabilities of distinct neuronal populations.

Furthermore, we aspire to integrate ’-omic’ data, tools, and results, with a particular emphasis on single-cell or single-nucleus RNA sequencing data, and the means to combine and analyze both data sources. Our overarching objective is to facilitate collaboration, knowledge sharing, and advancements in the field of spinal cord injury research through this comprehensive and continually expanding repository.

### Test Bench for Neuronal Identification Methods

In its maiden application within the NeuroCluedo repository, we have embarked on a comparative analysis of neuronal identification methods, namely manual, threshold-based, and neuronal network-based approaches. The purpose is to evaluate their potential and limitations. Within the repository, we have established a test bench that is accessible to all for conducting further comparisons. Our current comparisons underscore the superiority of the neuronal network method over the manual and threshold-based methods.

Traditional manual methods have been a prevalent choice for neuronal identifications, probably due to their simplicity and minimal technological requirements. However, our results suggest that manual methods are the most time-consuming and exhibit limited repeatability and reproducibility. The latter two drawbacks are of particular concern, given the substantial disparities observed, sometimes exceeding 100 neurons in repeated measurements of sections with fewer than 200 neurons. This leads to the observation that the most commonly identified neurons occur in only one out of every five analyses. Even with an increase in expertise from the analyst, our analyses reveal that experienced researchers struggle to consistently deliver repeatable identifications. Indeed, as a result of our study, our lab opted to duplicate the manual analyses for each image to enhance identification certainty.

The second method, reliant on thresholding to establish staining levels of neuronal and nuclear markers for neuron identification, represents an alternative approach (Reigada *et al*., 2015; Wang *et al*., 2019). The thresholding procedure here employed is semi-automated, requiring manual selection of the thresholding algorithm due to substantial variations in background and staining intensity across different preparations, and even within sections of the same preparation. Consequently, reproducibility issues may arise due to differences in the chosen thresholding algorithm. While this method offers time-saving benefits over manual approaches, its efficiency is constrained. Notably, the selected thresholds often lack specificity, resulting in the inclusion of objects that human observers do not identify as neurons. To address this issue, we integrated a manual selection of the gray matter within the macro to exclude objects identified outside of it (Reigada *et al*., 2015). Improvements in thresholding techniques may be achievable through advancements in image preparation, background correction, and the selection of thresholding algorithms, among other factors. However, the extent to which these efforts are justifiable remains a topic of debate, particularly given the promising potential of AI-based methods.

The neuronal network-based identification method we have tested excels in several aspects. It demonstrates exceptional specificity, effectively distinguishing autofluorescence and non-specific labeling from genuine neuronal staining. Notably, the neuronal network reduces the number of identified neurons inconsistent with manual analyses. While the initial time investment for training the neuronal network is substantial, the algorithm’s subsequent efficiency is evident as it can analyze hundreds of images within minutes without human intervention. Nevertheless, there are two notable limitations to consider. Firstly, it relies on Olympus software, which entails a costly license. Secondly, the neuronal network does not classify objects as neurons but assigns a probability to each pixel in the image, indicating the likelihood of it being part of a neuron. To overcome these limitations, researchers can explore open-source software solutions such as StarDist (Schmidt *et al*., 2018) or MIA (Körber, 2023). These open-source tools not only offer cost-effectiveness but are also well-suited for object identification. Furthermore, the flexibility to train the neuronal network under various conditions opens avenues for result improvement and application across diverse scenarios at minimal additional cost.

Although inconsistencies in neuron identifications emerged between observers and were particularly evident when compared to the semi-automatic macro initially employed in the Reigada *et al*. (2015) study, the accuracy analyses conducted suggest that the estimates of the total number of neurons in each section remain reasonably consistent across various methods and observers. The factors that reconcile the divergent identifications when calculating the total number of neurons fall outside the scope of this paper. However, it is essential to underscore that these factors affirm the validity of the results presented in the original paper by Reigada *et al*. (2015), despite the identification challenges associated with the method employed.

### Analysis of neuronal death and neuroprotection

In addition to exploring the potential for developing and testing image analysis tools, our objective was to leverage the stored images to investigate the distribution of neuronal death within the spinal cord in a mouse model of contusive SCI and identify the neurons protected by the anti-apoptotic drug ucf-101 treatment. We focused on sections located 0.6 to 1.2 mm caudal to the injury epicenter, as this region initially showed the most pronounced neuroprotective effects of ucf-101 (Reigada *et al*., 2015). For neuron identification, we employed the neuronal network developed in this study. To ensure data comparability across different individuals and experimental conditions, we utilized thin plate spline techniques for image registration, aligning them with reference sections from the Allen reference atlas for the mouse spinal cord. This approach facilitated not only the quantification of neurons but also their spatial localization and their allocation within Rexed laminae.

The findings from these analyses shed light on the distinctive pattern of neuronal death that unfolds the current model of SCI. Firstly, they confirm the existence of a gradient of neuronal death concerning the proximity to the injury epicenter, consistent with prior observations in Reigada *et al*. (2015). According to this pattern, loss of neurons is massive at distances below 0.6 mm from the epicenter. This effect gradually wanes, ultimately resulting in less than 50% neuronal loss at a distance of 1.2 mm. We observed a similar pattern of neuronal loss within 400 microns caudal to the injury epicenter, with subsequent progressive survival as the distance from the epicenter increased in a recent investigation, examining neuronal survival following SCI in both wild-type (WT) and XIAP overexpressing C57BL/6 mice (Reigada *et al*., 2023). In a similar contusive model with higher severity (75 *vs* 50 Kdynes), Sauerbeck *et al*. (2021) observed a substantial number of neurons preserved in sections located 600µm rostral to the injury epicenter in female mice. This contrasts with our current findings where neurons are nearly absent at this distance. These disparities could suggest that caudal preservation is more constrained than rostral preservation, a possibility supported by our results in Reigada *et al*. (2023) but not in the data that underpins the present study (Reigada *et al*., 2015).

Within the transverse section of the spinal cord, our results further suggest that cell death predominantly impacts neurons located in intermediate regions. Specifically, this phenomenon pertains to the nuclei and laminae situated adjacent to the ependyma, with a particular emphasis on laminae 5 and 10, as well as nuclei D and LDCom. To what extent the observed pattern reveals the intrinsic vulnerability of neuronal populations or differences in the deleterious stimuli they confront is out of the scope of the present study. It is important to acknowledge that the consistency of this pattern within the present sample and across various studies remains open to question. Individual variability appears to be substantial, and our current results allude to a general trend rather than a uniform observation.

Despite the extensive body of literature involving histological analyses of neuronal death in mouse models of SCI (a search on PubMed retrieves more than 400 articles on Spinal Cord injury AND neuronal death AND mouse), information on neuronal death distribution is almost lacking. In one of the few descriptions, Matsushita and colleagues (Matsushita *et al*., 2000) described the accumulation of TUNEL-positive neurons in the dorsal horns following ischemia/reperfusion SCI, in contrast to the accumulation of neuronal death in the intermediate region we have observed. While differences in the injury model may offer an explanation, the abundance of TUNEL-positive neurons in the most dorsal laminae may also reflect the higher density of neurons in this particular region.

In our 2015 study, we investigated the neuroprotective impact of ucf-101 in the context of SCI. Ucf-101 (5-[5-(2-nitrophenyl) furfuryliodine] 1,3-diphenyl-2-thiobarbituric acid) is a heterosynthetic cyclic compound known for its demonstrated neuroprotective effects in animal models of cerebral ischemia (Althaus *et al*., 2007) and other neurodegenerative diseases (Goffredo *et al*., 2005; Ding *et al*., 2009). According to our original study, the administration of ucf-101 resulted in a noteworthy increase in the survival of neurons located within the 0.6 - 1.2 mm caudal to the injury epicenter. Consistent with these findings, our current analysis corroborates ucf-101-induced neuroprotection in sections situated at 0.6 and 0.8 mm caudal from the injury epicenter. However, we have not detected significant neuroprotection at greater distances. These discrepancies arise from the advances in the methodology for neuron identification, particularly notable for sections located 1.0 mm caudal from the epicenter. In these sections, the threshold-based method employed in the original article identified multiple artifacts or background signal as neurons, which were excluded by the neuronal network-based method. Nonetheless, it’s worth noting that the limited availability of samples with sufficient preservation for analysis at 1.2 mm may also contribute to the observed differences.

Our image analysis also allowed us to pinpoint specific Rexed laminae and nuclei that exhibit enhanced neuroprotection. In particular, we have observed increased survival of interneurons within laminae L4, L5l, L5m, L7, and the D nucleus. Interestingly, the increased neuroprotection within the intermediate region of the transverse section broadly agrees with the region of higher neuronal death following injury. Collectively, the spatial distribution of surviving neurons suggests that the protective effects of ucf-101 are primarily confined to neurons under strong deleterious pressures, particularly those closest to the injury epicenter. Considering the specific effects of ucf-101 on the intrinsic pathway of apoptosis, this observation could be interpreted as an indication that this pathway is active in the border of spared tissue surrounding the injury epicenter caudal to the injury. This observation aligns with previous clinical data and animal models of SCI (see Lu *et al*., 2000, and references therein) and related conditions such as stroke (Fricker *et al*., 2018).

The patterns of spatial distribution of neuronal death after SCI and the neuroprotective effects of ucf-101 described here can be considered, at best, as hypotheses to be tested. Variability among individuals and sections makes it difficult to draw more consistent patterns. Whereas a substantial part of the observed variability arises from biological or injury differences among individuals and constitutes a key information to characterize the responses to injury and treatment, variability is also a consequence of histological artifacts. For example, we observed a large difference in the number of neurons identified in 3 sections from one control individual (216, 260, and 503 neurons), which was associated with a high staining intensity in one of the sections. To address variability in neuronal counts within conditions, we have employed the median as a centrality measurement through most analyses. However, further analysis and quality controls may be required to deal with histological artifacts. On the contrary, dealing with biological variability will benefit from increasing the sample and incorporating additional information on the injury. Our intention for future analyses is to incorporate samples from additional studies carried out by the group (Muñoz de Galdeano *et al*., 2018; Reigada *et al*, 2023) as well as to additional sources of information (*eg*. tissue sparing) so we can characterize the variability of neuronal death through individuals and studies.

Several steps of the present image analyses can be improved. The first one concerns image registration, the process of overlaying two or more images to achieve maximum correspondence, which we carried out using BigWarp, a tool for landmark-based deformable image alignment (Bogovic *et al*., 2016). A number of image registration methods have been proposed, including several specifically developed for histological images (for example, cWR; Wang *et al*., 2014). However, in our hands, neither this nor more general tools such as pyStackReg (Thevenaz *et al.,* 1998) or Elastix (Marstal *et al*., 2016), yielded better results that BigWarp (data not shown). Morphological deformations together with staining differences make automatic registration of histological sections highly challenging. Indeed, a recent challenge demonstrated that image registration of histological sections requires the development of specific tools which complicates or even make it impossible for regular users to apply those (Borovec *et al*., 2020). In all, results from image registration using BigWarp can be considered optimal, although several aspects should be improved. On the one hand, landmarks employed for thin plate spline deformations were not standardized, but both their number and position were selected subjectively to obtain the best alignment between the section under study and the Allen’s reference. On average, we employed 35 landmarks but the number varied upon the preservation of the spine, so that well-preserved gray substances required few landmarks (down to a minimum of 11) while highly deformed ones required up to 53 landmarks to obtain an acceptable alignment. Moreover, difficulties in the identification of homologous positions of landmarks in some damaged sections also made deformations (and therefore the subsequent quantifications) questionable. Future studies should focus on defining a set of identifiable landmarks warranting homologous positions in the reference and moving image. On the other hand, we employed the Allen’s reference as the moving image that was aligned to the image under analysis. While this approach ensures that the section and its component neurons are not deformed, it forced manual counting of the neurons in each nucleus and lamina. This step can be automated if we employ the spinal cord section under analysis as the moving image (instead of the reference map), so that the regions of interest (ROIs, i.e laminae and nuclei) defined in the reference image can be automatically quantified. Recently Fiederling *et al*. (2021) have described procedures for neuronal quatification based on registering to 3D reference volumes derived from the Allen Spinal Cord Atlas (Lein *et al*., 2007) instead to the segment 2D annotations of this atlas. In future studies we will test whether this approach solves the registration problems we have identified.

A final limitation of our methods concerns the quantification of neurons in each Rexed lamina or spinal nucleus. Even though this regionalization has well-established biological roots, each region having specific features in terms of neuronal function, chemistry, and morphology, this way of analyzing the spatial distribution of the surviving neurons has limitations. One the one hand, the boundaries of laminae and nuclei are section specific and, therefore, the assignment of neurons, in cases such as the intermediolateral nucleus, can be questionable when using Allen’s reference that is derived from a different section of a different individual. On the other hand, the use of predefined regions restricts the spatial changes in neuronal death to those detectable at this regional scale, i.e., patterns of neuronal survival/cell death unrelated to the proposed regionalization are lost or at least diluted. In this respect, we consider that direct analysis of the spatial location of each surviving neuron through spatial statistics approaches will be more suitable for detecting patterns or correlations of neuronal death within the spinal cord as well as for reducing subjectivity. In this respect, the procedures employed in the Blue Brain project (Erö *et al*., 2018), by defining the voxels in a reference volume as the basic units for quantifying neuronal abundance, can be a valid alternative.

## CONCLUSIONS AND PERSPECTIVES

In the last decades, there has been significant efforts aimed at repairing SCI. Our understanding of SCI pathophysiology has expanded and multiple therapeutic treatments have been developed, showing varying degrees of success in animal models. Despite these advancements, we have yet to discover a single therapy that demonstrates both efficacy and safety in clinical trials, qualifying it for clinical practice. This study not only contributes to the understanding of SCI but also aligns with an exciting area where the integration of advanced techniques, such as 3D mapping of neurons based on imagen analysis and the revolutionary Single Cell RNA sequencing, is redefining the boundaries of neuroscience research. Leading initiatives, such as the Blue Brain Project, have been at the forefront of this movement. The combined analysis of spinal cord images with single-cell RNA sequencing data opens new avenues for addressing fundamental questions in neurobiology and lays the groundwork for future clinical applications. However, there is a noticeable absence of information concerning the spatial distribution of the different forms of neuronal death over time. Available spatial information dates back to 90s and early 2000s and is restricted to necrotic and apoptotic cell deaths, particularly in rat models of SCI (see, for example, Liu *et al*., 1997). On the contrary, thousands of histological preparations from mouse models of SCI are employed, in all but a few articles, to illustrate differences among conditions or to compare specific aspects of these conditions. In light of ethical imperatives to minimize the use of laboratory animals, we have carried out a proof-of-concept to explore the potential of leveraging recent technological advancements to maximize the wealth of information derived from the histological data gathered through the years. Our results, available at NeuroCLUEDO repository, confirm that previous histologies preserve information key to decode and understand the processes responsible for the spatial distribution of neuronal death and the neuroprotective effects of therapies under development.

## ACKNOWLEDGMENTS

This study and the NeuroCLUEDO project has been co-financed by the European Union (FEDER) “A way to make Europe”, We thank for their technical and logistic support to the Fundación del Hospital Nacional de Parapléjicos para la Investigación y la Integración (FUHNPAIIN) and the microscopy facility of the Experimental Neurology Unit, Hospital Nacional de Prapléjicos, Toledo.

