## Supplementary material for "Reanalysis of published histological data can help to characterize neuronal death after Spinal Cord Injury": Suplementary Materials

I.Neuron identification methods

- Manual identification

Manual analysis of the 20 images comparison set was carried out by six observers, each of them analyzing twice 6 images from the set except for the NIB who analyzed all images once. Images were provided to the observer for analysis on a daily basis with repeated analysis of the same image separated at least 7 days. Observers were not informed of image analysis repetition. The distribution of the images among observers is shown in Table S1. The number and position of neurons in each image were analyzed using the Cell Counter plugin from Fiji and stored in .xml files. The time spent analyzing each image was also recorded. Only those cells double-stained against NeuN and DAPI located within the gray substance were taken into consideration. According to their skills to identify neurons in the damaged spinal cord, observers were classified as Beginners (OB141 & NIB, no previous experience in analyzing neurons, n=2), Advanced (OB109 & OB168, experience using image software to analyze neurons from tissue sections and cell cultures, n=2), and Expert (OB132 & OB 180, experience using image software to analyze neurons in injured spinal cord, n=2). All observers were provided with specific instructions, available at https://osf.io/snjwk/ together with further details on the distribution of images among observers and the obtained .xml files.

- Threshold-based identification

We employed the original results published by Reigada *et al*. (2015) together with additional unpublished data. In these analyses, the authors employed a custom Fiji macro (Threshold macro.txt, available at https://osf.io/7agyq/) that uses basic processing techniques (background subtraction, thresholding, watershed, etc.) to segment the images and automatically detect cell nuclei and NeuN staining. The resulting masks of identified neurons were exported as .tiff files available at NeuroCLUEDO (Threshold identifications.zip, available at https://osf.io/7agyq/) together with the ImageJ macro.

- Neural network-based identifications

To build up the neural network for neuronal identification we employed the deep-learning-based image analysis approach TruAI integrated CellSens Dimensions software (Olympus), which uses deep convolutional neural network architecture for object segmentation. We trained the neural network with 995 manually segmented neurons from 25 images of the training set (ground truth). Neurons were selected from all across the gray matter of control and injured spinal cords. The background surrounding each neuron as well as artifacts were also identified. The training was carried out using the Deep Learning module operating under the Standard Network configuration, which utilizes a U-Net architecture with 32 feature maps in the first convolution layer, on 300000 iterations with 5 checkpoints every 60000 iterations. Validation was carried out on 10% of the segmented images (not employed for training) in each iteration and expressed as a percentage of correctly identified neurons. CellSens Dimensions software determines the accuracy of TruAI object segmentation predictions compared with the manually identified neurons in single images. A minimum of 70% congruence between manual and neural network identifications was set. The resulting neural network assigns a probability of being part of a neuronal nucleus to every pixel in the image. To identify neuronal nuclei, we considered only those particles composed of pixels above 50% probability and an area above 25 $\mu m^2$. The designed neural network (NC-AI1), as well as all identifications and every code employed in these identifications, are available at NeuroCluedo (https://osf.io/7agyq/).

### II. Data analysis

- Comparison of neuronal identifications

The neurons identified by each method were compared by overlapping their positions or segmentations obtained with each method for each image. Overlapping of the manual identifications was used to establish the consistency among repeated identifications by the same observer and between observers (Table S1). To evaluate the identifications provided by the thresholding and neuronal network-based methods, each manually identified neuron in a given image was first characterized by the ratio of the number of times observers identified it divided by the total number of times the image was analyzed. The resulting ratios were used to define the reliability of each identified neuron and employed as ground truth to analyze which neurons were identified by each method.

- Repeatability and reproducibility of neuronal quantifications

The repeatability and reproducibility of the estimated total number of neurons were analyzed using the *difference plot* created by Bland and Altman (1986). We also employed linear models to analyze the effect of the number of neurons, section features, or the observer experience on repeatability and reproducibility. Models were adjusted using the *lm* function and later compared using *anova* function of R (R Core Team, 2017). Repeatability and reproducibility of neural network identifications, as well as repeatability of threshold-based identifications, were assumed as 100%.

- Accuracy of neuronal quantifications

Accuracy was quantified by comparing the neurons identified with each method with the reference neurons for each image. Given that the actual number of neurons captured in each section is unknown, we set the reference number of neurons (RNN) per section from the weighted consensus of the manual estimations for each image. RNN in each section was calculated by means of the probability of each identified neuron to really correspond to an actual neuron, so that neurons identified in all analyses of an image (3 out of 3, or 5 out of 5) are assigned a probability of 100%, those identified by 4 out of 5 observers are assigned 80% (100*4/5), 2 out of 3 are assigned a 66% (100*2/3) and so on. Then, the total number of neurons is estimated according to the following formula:

- For sections analyzed 5 times:

  Total Nº of neurons = Nº of neurons 100% (5/5) + Nº of neurons 80% (4/5) x 0.8 + Nº of neurons 60% (3/5) x Nº of neurons 40% (2/5) x 0.4 + Nº of neurons 20% (1/5) x 0.2.

- For sections analyzed in 3 replicates:

  Total Nº of neurons = Nº of neurons 100% (3/3) + Nº of neurons 66% (2/3) x 0.66 + Nº of neurons 33% (1/3) x 0.33.

The total number of neurons identified by each observer or method were compared to the RNN graphically and through Pearson correlation.

III. Reanalysis of ucf-101 effects on neuronal survival

- Images

We focused this analysis on the caudal penumbra sections (0.6-1.2 mm from the injury epicenter) where significant differences in the number of surviving neurons were identified between vehicle and ucf-101 treated animals in Reigada *et al*. (2015). A total of 42 sections were included in the analyses, distributed as detailed in Table S2.

- Neuronal detection

Raw confocal images in .lif format were converted into individual .tif files using Fiji. Processing involved the maximum intensity projection of the focal planes of the confocal image and its conversion into RGB format. Neurons were identified using the NC-AI1 neuronal network developed in section B.4. NC-AI1 assigns to every pixel in the image a probability of being part of a neuronal nucleus. Those particles above 25 $\mu m^2$ composed of pixels above 50% probability were classified as neuronal nuclei. NC-AI1 identifications and post-processed images are available at https://osf.io/fhxqs/.

- Image Registration and Quantification of Neurons in Rexed laminae

Images from each section under analysis were registered to a reference map of the Rexed laminae in thoracic segment 11 obtained from the Mouse Spinal Cord section of the Allen Brain Atlas (https://mousespinal.brain-map.org/imageseries/showref.html, last date accessed august 2023) to obtain an estimation of the neurons present in each lamina and nucleus comprised in this atlas. To do so, we employed the BigWarp tool (Bogovic *et al*., 2016) from Fiji, setting the image with identified neurons as the target image and the atlas image as the moving one (Figure S1). BigWarp carries out a non-affine Thin Plate Spline-based deformation of the moving image to adjust its landmarks to the corresponding ones in the target image. Landmarks and registered images are available at https://osf.io/fhxqs/.

- Neuronal quantification, data processing, and data analysis

The number of neurons in each lamina and nucleus was established by two observers (PRA and MND) who manually counted all sections. Overlapping of the atlas on the processed images (including neuronal identifications) was employed to quantify the number of neurons in each lamina and nucleus. In case of disparity among the observers, the image was reanalyzed and discussed until a consensus was reached. Raw data were recorded in an Excel matrix detailing for each section the left, right, and mean values of the number of neurons in each lamina and nuclei except for lamina 10 and lumbar dorsal commissural nucleus (LDCom) which have unique values. Raw data were processed to retain only the mean values while incorporating variables on the individual, the image and section codes, the treatment/condition, and the section position relative to the injury epicenter. Matrices are available at https://osf.io/fhxqs/. Data were analyzed using Social Science Statistics online tool (https://www.socscistatistics.com/tests/mannwhitney/) for Mann-Whitney U statistical testing and Auto-Weka (Thornton *et al*., 2013) for data classification.

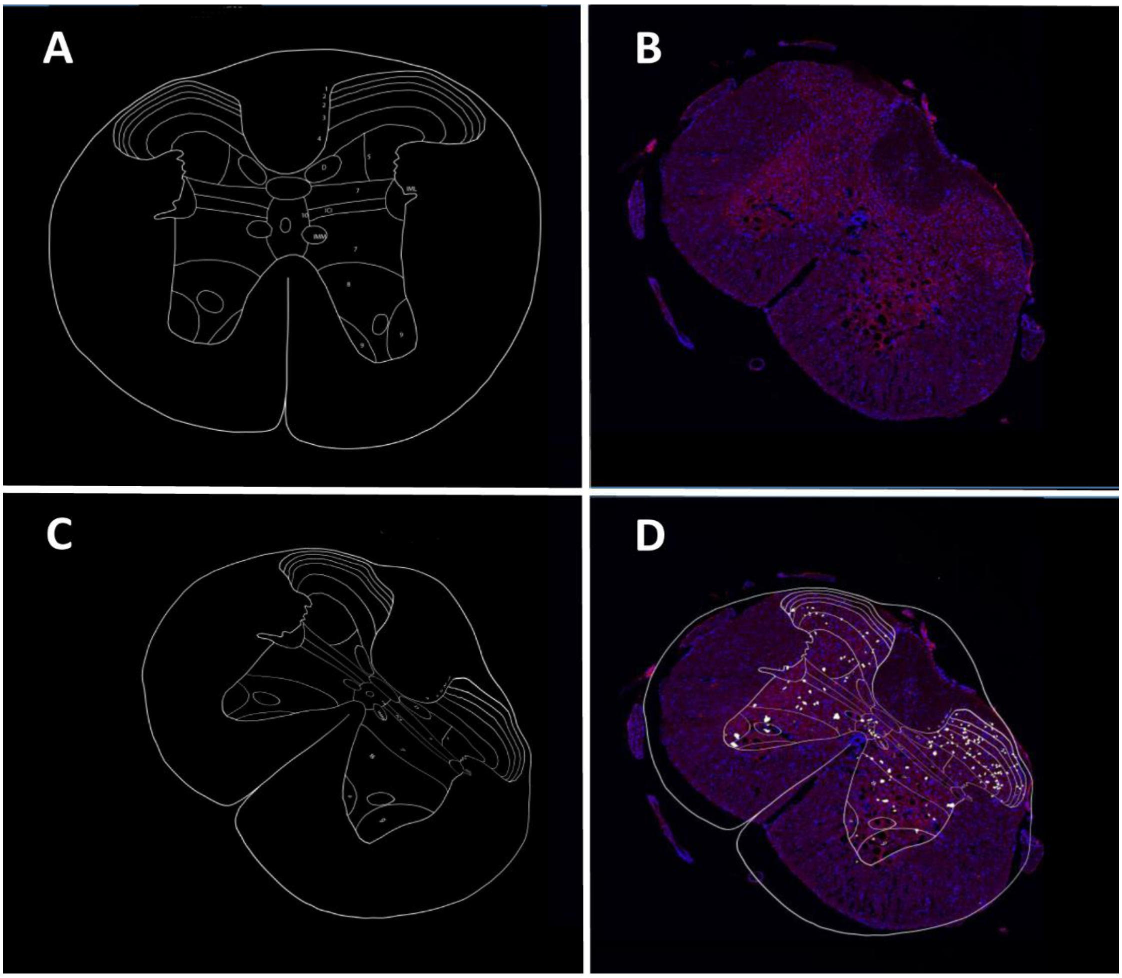


**Figure S1. Image registration**. A) Atlas of mice T11 transversal section employed as moving image (modified from Allen Brain Atlas, https://mousespinal.brain-map.org/imageseries/showref.html, last date accessed August 2023). B) Stained image employed as target section. This image comprises DAPI and NeuN layers plus the Neuronal Network-based identifications. C) Deformation of the moving image (atlas) to match the stained section form D). Deformations were focused on matching the gray matter contour, without adjusting for the spinal cord contour.

| Image | Treatment | Observer | | | | | | Nº of analyses |
|---|---|---|---|---|---|---|---|---|
| | | OB180 | OB132 | OB141 | OB109 | OB168 | NIB | |
| 175 UCF RGB.tif | Ucf-101 | | 2 | | | | 1 | 3 |
| 176 UCF RGB.tif | Ucf-101 | | | | 2 | | 1 | 3 |

| | | | | | | | | |
|---|---|---|---|---|---|---|---|---|
| 177 UCF RGB.tif | Ucf-101 | 2 | | 2 | | | 1 | 5 |
| 178 UCF RGB.tif | Ucf-101 | 2 | | | | 2 | 1 | 5 |
| 179 UCF RGB.tif | Ucf-101 | | | | 2 | 2 | 1 | 5 |
| 180 UCF RGB.tif | Ucf-101 | | 2 | | 2 | | 1 | 5 |
| 181 UCF RGB.tif | Ucf-101 | | | 2 | | | 1 | 3 |
| 182 UCF RGB.tif | Ucf-101 | | | | | 2 | 1 | 3 |
| 183 UCF RGB.tif | Ucf-101 | 2 | 2 | | | | 1 | 5 |
| 184 UCF RGB.tif | Vehicle | | | 2 | 2 | | 1 | 5 |
| 185 UCF RGB.tif | Vehicle | 2 | | | 2 | | 1 | 5 |
| 186 UCF RGB.tif | Vehicle | | 2 | 2 | | | 1 | 5 |
| 187 UCF RGB.tif | Vehicle | | | | | 2 | 1 | 3 |
| 188 UCF RGB.tif | Vehicle | | | 2 | | 2 | 1 | 5 |
| 189 UCF RGB.tif | Vehicle | | 2 | | | | 1 | 3 |
| 190 UCF RGB.tif | Vehicle | 2 | | | | | 1 | 3 |
| 191 UCF RGB.tif | Control | 2 | | | | | 1 | 3 |
| 193 UCF RGB.tif | Control | | | 2 | | | 1 | 3 |
| 195 UCF RGB.tif | Control | | 2 | | | 2 | 1 | 5 |
| 197 UCF RGB.tif | Control | | | | 2 | | 1 | 3 |

**Table S1. Design for the analysis of the manual identification of neurons.** The table details for each image, its code, the type of treatment, the observers who analyzed it, the number of replications performed by each observer, and the total number of analyses performed on each spine section. The observers are indicated by a code (OB + random number) to preserve their anonymity. NIB corresponds to experiment coordinator Nadia Ibañez Barranco.

| **Condition** | **Control** | **Vehicle** | | | | **UCF-101** | | | |
|---|---|---|---|---|---|---|---|---|---|
| $d_{epicenter}$ (mm) | - | 0.6 | 0.8 | 1.0 | 1.2 | 0.6 | 0.8 | 1.0 | 1.2 |
| N. of images | 6 | 5 | 5 | 6 | 5 | 5 | 5 | 4 | 1 |

**Table S2**. **Number of images** (sections) analyzed in each condition and distance to the epicenter. Additional data on the samples are available at https://osf.io/fhxqs/.

| **Image/section** | **Condition** | **Number of analysis** | **Overlap** | | | | |
|---|---|---|---|---|---|---|---|
| | | | **1** | **2** | **3** | **4** | **5** |
| 175 UCF RGB.tif | Ucf-101 | 3 | 105 | 104 | 283 | | |

| | | | | | | | |
|---|---|---|---|---|---|---|---|
| 176 UCF RGB.tif | Ucf-101 | 3 | 298 | 106 | 87 | | |
| 177 UCF RGB.tif | Ucf-101 | 5 | 56 | 42 | 46 | 67 | 218 |
| 178 UCF RGB.tif | Ucf-101 | 5 | 78 | 44 | 64 | 72 | 230 |
| 179 UCF RGB.tif | Ucf-101 | 5 | 43 | 20 | 37 | 58 | 131 |
| 180 UCF RGB.tif | Ucf-101 | 5 | 87 | 20 | 7 | 4 | 11 |
| 181 UCF RGB.tif | Ucf-101 | 3 | 40 | 25 | 29 | | |
| 182 UCF RGB.tif | Ucf-101 | 3 | 55 | 49 | 159 | | |
| 183 UCF RGB.tif | Vehicle | 5 | 82 | 39 | 19 | 19 | 7 |
| 184 UCF RGB.tif | Vehicle | 5 | 198 | 95 | 16 | 1 | 0 |
| 185 UCF RGB.tif | Vehicle | 5 | 30 | 11 | 0 | 0 | 0 |
| 186 UCF RGB.tif | Vehicle | 5 | 40 | 11 | 0 | 0 | 0 |
| 187 UCF RGB.tif | Vehicle | 3 | 15 | 5 | 0 | | |
| 188 UCF RGB.tif | Vehicle | 5 | 93 | 24 | 14 | 6 | 3 |
| 189 UCF RGB.tif | Vehicle | 3 | 263 | 89 | 85 | | |
| 190 UCF RGB.tif | Vehicle | 3 | 37 | 7 | 3 | | |
| 191 UCF RGB.tif | Control | 3 | 171 | 161 | 216 | | |
| 193 UCF RGB.tif | Control | 3 | 163 | 150 | 427 | | |
| 195 UCF RGB.tif | Control | 5 | 167 | 106 | 110 | 105 | 196 |
| 197 UCF RGB.tif | Control | 3 | 228 | 124 | 224 | | |

**Table S3. Coherence in manual identifications**. The table shows the results obtained after overlapping the manual identifications for each image of the twenty spinal cord sections. The table details the code of each image/section, the condition of the individual, the total number of analyses performed on each section, whereas Overlap describes the number of neurons with overlapped counts in each section. The value of 1 refers to neurons that have only been identified in one analysis, the value of 2 corresponds to neurons identified in two analyses, and so on until the values of 5.

| | | | Number of counts | | | | | | |
|---|---|---|---|---|---|---|---|---|---|
| **Image** | **Treatment** | **Analysis** | **0** | **1** | **2** | **3** | **4** | **5** | **Total** |
| 178 UCF RGB.tif | Ucf-101 | Macro + | 33 | 11 | 7 | 17 | 29 | 160 | 257 |
| | | Macro - | | 66 | 35 | 42 | 44 | 59 | |
| | | IA + | 64 | 57 | 39 | 50 | 70 | 227 | 507 |
| | | IA - | | 13 | 3 | 7 | 1 | 1 | |
| 180 UCF RGB.tif | Ucf-101 | Macro + | 83 | 9 | 9 | 5 | 2 | 11 | 119 |
| | | Macro - | | 73 | 11 | 0 | 2 | 0 | |
| | | IA + | 27 | 8 | 8 | 5 | 3 | 10 | 61 |
| | | IA - | | 76 | 12 | 1 | 1 | 0 | |
| 183 UCF RGB.tif | Vehicle | Macro + | 220 | 28 | 21 | 15 | 11 | 10 | 305 |
| | | Macro - | | 59 | 13 | 4 | 8 | 0 | |
| | | IA + | 42 | 18 | 18 | 11 | 13 | 7 | 109 |
| | | IA - | | 72 | 18 | 7 | 6 | 0 | |
| 188 UCF RGB.tif | Vehicle | Macro + | 252 | 55 | 17 | 8 | 6 | 3 | 341 |
| | | Macro - | | 25 | 4 | 4 | 0 | 0 | |
| | | IA + | 35 | 9 | 8 | 6 | 4 | 3 | 65 |
| | | IA - | | 80 | 15 | 7 | 2 | 0 | |
| 195 UCF RGB.tif | Control | Macro + | 152 | 44 | 45 | 71 | 75 | 173 | 560 |
| | | Macro - | | 119 | 48 | 32 | 22 | 19 | |
| | | IA + | 55 | 70 | 71 | 79 | 81 | 188 | 544 |
| | | IA - | | 82 | 26 | 17 | 17 | 4 | |

**Table S4. Agreement among neuronal identifications**. The table shows the agreement between neuronal identifications obtained using manual, threshold-based (macro), and artificial intelligence-based (AI) methods. The table details the code of each image (Image), the condition (treatment), the employed identification method (Analysis), and the number of identified neurons (Number of counts). The Number of counts is split by the number of neurons identified in 0 to 5 manual analyses. The value zero corresponds to neurons only identified by threshold or artificial intelligence methods. Value 1 refers to neurons identified in a single manual analysis and the values 2, 3, 4, and 5 are neurons that have been identified in two, three, four, and five manual analyses, respectively. “Macro +” and “IA +” indicate the number of neurons identified by each method, while macro - and IA - refer to neurons that have only been identified by manual analysis. Total indicates the total number of neurons identified in at least one manual analysis.

| **Image** | VOLUNTEERS | | | | | |
|---|---|---|---|---|---|---|
| | **OB168** | **OB141** | **OB109** | **OB132** | **OB180** | **OB184** |
| 175 UCF RGB.tif | | | | 375/26 min<br>487/20 min | | 431/39 min |

| | | | | | | |
|---|---|---|---|---|---|---|
| 176 UCF RGB.tif | | | 202/25 min<br>138/17 min | | | 244/21 min |
| 177 UCF RGB.tif | | 399/25 min<br>294/20 min | | | 344/29 min<br>382/11 min | 327/17 min |
| 178 UCF RGB.tif | 352/25 min<br>363/16 min | | | | 345/14 min<br>398/15 min | 352/20 min |
| 179 UCF RGB.tif | 216/17 min<br>264/8 min | | 214/13 min<br>191/14 min | | | 222/14 min |
| 180 UCF RGB.tif | | | 42/9 min<br>29/7 min | 77/11 min<br>113/20 min | | 21/4 min |
| 181 UCF RGB.tif | | 53/13 min<br>92/30 min | | | | 71/15 min |
| 182 UCF RGb.tif | 206/14 min<br>231/13 min | | | | | 224/21 min |
| 183 UCF RGB.tif | | | | 44/18 min<br>115/20 min | 54/10 min<br>65/8 min | 51/11 min |
| 184 UCF RGB.tif | | 104/28 min<br>232/32 min | 40/16 min<br>16/5 min | | | 179/41 min |
| 185 UCF RGB.tif | | | 28/6 min<br>17/10 min | | 2/5 min<br>0/2 min | 6/9 min |
| 186 UCF RGB.tif | | 0/2 min<br>0/2 min | | 21/18 min<br>77/20 min | | 0/39 min |
| 187 UCF RGB.tif | 0/3 min<br>18/5 min | | | | | 0/5 min |
| 188 UCF RGB.tif | 53/14 min<br>31/7 min | 69/12 min<br>118/20 min | | | | 15/6 min |
| 189 UCF RGB.tif | | | | 241/20 min<br>362/17 min | | 145/29 min |
| 190 UCF RGB.tif | | | | | 20/8 min<br>22/10 min | 16/9 min |
| 191 UCF RGB.tif | | | | | 276/12 min<br>365/9 min | 507/67 min |
| 193 UCF RGB.tif | | 690/60 min<br>504/36 min | | | | 605/39 min |
| 195 UCF RGB.tif | 352/22 min<br>454/21 min | | | 476/29 min<br>379/15 min | | 542/32 min |
| 197 UCF RGB.tif | | | 282/29 min<br>320/21 min | | | 568/31 min |

**Table S5: Number of neurons identified in the spinal cord sections.** For each analyst, the number of identified neurons and the time (in minutes) spent in the analysis of each image is detailed. The code OB168, etc. anonymously identifies the analyst involved. Repeated measurements for an image and observer are shown except for observer OB184.

| | | Analysis Method | |
|---|---|---|---|
| **Image** | **RNN** | **Macro** | **AI** |
| 175 UCF RGB.tif | 387 | 336 | 311 |
| 176 UCF RGB.tif | 257 | 317 | 305 |
| 177 UCF RGB.tif | 327 | 364 | 416 |
| 178 UCF RGB.tif | 359 | 195 | 466 |

| | | | |
|---|---|---|---|
| 179 UCF RGB.tif | 216 | 0 | 331 |
| 180 UCF RGB.tif | 44 | 48 | 42 |
| 181 UCF RGB.tif | 59 | 50 | 117 |
| 182 UCF RGB.tif | 210 | 252 | 275 |
| 183 UCF RGB.tif | 63 | 116 | 103 |
| 184 UCF RGB.tif | 88 | 50 | 133 |
| 185 UCF RGB.tif | 10 | 31 | 18 |
| 186 UCF RGB.tif | 12 | 0 | 0 |
| 187 UCF RGB.tif | 8 | 0 | 0 |
| 188 UCF RGB.tif | 44 | 122 | 51 |
| 189 UCF RGB.tif | 232 | 103 | 118 |
| 190 UCF RGB.tif | 20 | 130 | 35 |
| 191 UCF RGB.tif | 380 | 326 | 260 |
| 193 UCF RGB.tif | 581 | 500 | 590 |
| 195 UCF RGB.tif | 422 | 387 | 505 |
| 197 UCF RGB.tif | 383 | 434 | 548 |

**Table S6. Total number of neurons in each section.** The table shows the Reference Number of Neurons (RNN) as well as the estimates from the threshold-based and Neuronal Network-based methods for each image under analysis.

| Condition & distance to injury epicenter | Number of sections |
|---|---|
| Control | 4 |
| Vehicle, 0.6 µm | 4 |
| Vehicle, 0.8 µm | 4 |
| Vehicle, 1.0 µm | 6 |
| Vehicle, 1.2 µm | 6 |
| UCF-101, 0.6 µm | 5 |
| UCF-101, 0.8 µm | 5 |
| UCF-101, 1.0 µm | 4 |
| UCF-101, 1.2 µm | 1 |

**Table S7. Number of images** for each condition and distance to the epicenter.

| | | L1 | L2O | L2I | L3 | L4 | L5L | L5M | D | L7 | ICI | IML | L7B | IMM | L8 | L9 | L10 | LDC | Total |
|---|---|---|---|---|---|---|---|---|---|---|---|---|---|---|---|---|---|---|---|
| Control | # | 26 | 23 | 22 | 50 | 37 | 20 | 8 | 4 | 11 | 9 | 3 | 28 | 3 | 17 | 4 | 19 | 11 | 283 |
| | % | 9 | 8 | 8 | 18 | 13 | 7 | 3 | 1 | 4 | 3 | 1 | 10 | 1 | 6 | 1 | 7 | 4 | 100 |
| Allen Atlas | # | 15 | 48 | 59 | 126 | 140 | 103 | 24 | 7 | 31 | 28 | 13 | 51 | 4 | 46 | 16 | 37 | 30 | 778 |
| | % | 2 | 6 | 8 | 16 | 18 | 13 | 3 | 1 | 4 | 4 | 2 | 7 | 1 | 6 | 2 | 5 | 4 | 100 |
| ratio %Ctrl/ %Allen | | 4.8 | 1.3 | 1.0 | 1.1 | 0.7 | 0.5 | 0.9 | 0.6 | 1.0 | 0.9 | 0.6 | 1.5 | 2.1 | 1.0 | 0.7 | 1.4 | 1.0 | |
| ratios above 2 fold are marked in red | | | | | | | | | | | | | | | | | | | |

**Table S8. Median number of neurons per Rexed laminae in controls and the Allen Atlas of adult mouse spinal cord (T11).** Percentage relative to the total number of neurons in a section is also shown to evaluate the agreement between data from the Allen reference atlas and our results. The last row shows the ratio between the percentages of neurons in each laminae estimated from the Allen atlas and our data. Major changes (above 2 fold) are shown in red type.

| Treatment | D.T.E. | L1* | L2O | L2I* | L3* | L4 | L5L* | L5M | D | L7 | ICI* | IML | L7B* | IMM** | L8** | L9 | L10 | LDC | Total |
|---|---|---|---|---|---|---|---|---|---|---|---|---|---|---|---|---|---|---|---|
| Control | - | 26 | 23.0 | 22 | 50.0 | 37.0 | 20 | 8.0 | 4.0 | 10 | 8 | 3 | 28 | 3 | 17 | 4 | 19.0 | 10 | 292 |
| Vehicle | 0.6 | 0** | 0** | 0** | 2** | 2** | 0** | 0* | 0 | 0** | 0** | 0* | 2** | 0** | 1** | 0** | 0* | 0** | 7** |
| | 0.8 | 2** | 5* | 7** | 16** | 10** | 4* | 1* | 0 | 2* | 1** | 0* | 5** | 0** | 4* | 0* | 2* | 0** | 59** |
| | 1.0 | 9** | 7** | 9* | 20** | 9** | 6** | 2* | 1 | 3* | 2* | 3 | 12** | 1* | 6* | 2* | 6* | 2** | 100** |
| | 1.2 | 16* | 17 | 16 | 30* | 16 | 5 | 2 | 0 | 5 | 4 | 3 | 14 | 2* | 6 | 2 | 7 | 4 | 149 |
| Ucf-101 | 0.6 | 5 | 6 | 2 | 8 | 7 | 3 | 1 | 1 | 2 | 1 | 1 | 5 | 1 | 2 | 1 | 0 | 2 | 45 |
| | 0.8 | 7 | 7 | 6 | 10 | 14 | 10 | 1 | 2 | 5 | 2 | 2 | 6 | 0 | 5 | 1 | 5 | 1 | 80 |
| | 1.0 | 9 | 7 | 10 | 18 | 21 | 10 | 3 | 2 | 5 | 3 | 1 | 9 | 1 | 4 | 2 | 4 | 2 | 110 |
| | 1.2 | 16 | 7 | 7 | 14 | 19 | 18 | 4 | 4 | 6 | 2 | 3 | 5 | 1 | 7 | 4 | 2 | 3 | 119 |

**Table S9. Number of neurons in sections from un-damaged spinal cords and from damaged sampled 21 days after injury at 0.6, 0.8, 1.0, and 1.2 mm caudal to the contusion epicenter.** The table details the median values of the number of neurons in each section and in each laminae and nucleus of the T11 spinal cord segment before and after SCI with or without ucf–101 treatment. For vehicle data, statistically significant neuronal losses relative to control values are marked for each region and distance to the epicenter. For Ucf-101 data, protection (reduction in neuronal losses) is compared to the corresponding laminae and distance of the vehicle-treated individuals. * and ** indicates p<0.05 and p<0.01 significance after one-tailed Kruskal Wallies test and Dunn's posthoc test. A detailed description of the statistical results is available in OSF project (https://osf.io/n32z9/). D.T.E. indicates Distance to Epicenter, expressed in mm.
